# Analysis of Genomes of Bacterial Isolates from Lameness Outbreaks in Broilers

**DOI:** 10.1101/2020.11.04.369157

**Authors:** N. Simon Ekesi, Beata Dolka, Adnan A.K. Alrubaye, Douglas D. Rhoads

## Abstract

We investigated lameness outbreaks at commercial broiler farms in Arkansas. From Bacterial Chondronecrosis with Osteomyelitis (BCO) lesions, we obtained different isolates of distinct bacterial species. Genome assemblies for *Escherichia coli* and *Staphylococcus aureus* isolates show that BCO-lameness pathogens on farms can differ significantly. Genomes assembled from *Escherichia coli* isolates from three different farms were quite different from each other, and more similar to isolates from different hosts and geographical locations. The S aureus genomes were closely related to chicken isolates from Europe, and appear to have been restricted to chicken hosts for more than 40 years. Detailed analyses of genomes from this clade of chicken isolates with a sister clade of human isolates, suggests the acquisition of a particular pathogenicity island in the transition from human to chicken pathogen and that pathogenesis in chickens may depend on this mobile element. Phylogenomics is consistent with more frequent host shifts for *E. coli*, while *S. aureus* appears to be highly host restricted. Isolate-specific genome characterizations will help further our understanding of the disease mechanisms and spread of BCO-lameness, a significant animal welfare issue.

**Importance:** Detailed inspection of the genome sequences of different bacterial species associated with causing lameness in broiler chickens reveals that one species, *E. coli*, appears to easily switch hosts from humans to chickens and other host species. Conversely, isolates of *S. aureus* appear to be restricted to specific hosts. One potential mobile DNA element has been identified that may be critical for causing disease in chickens for *S. aureus*.

## Introduction

Lameness poses animal health welfare issues which results in significant losses in poultry production. Modern broilers selectively bred for rapid growth are particularly prone to leg problems (1). Bacterial chondronecrosis with osteomyelitis (BCO) is the leading cause of lameness in broiler and broiler breeder flocks (1–3). In birds that develop lameness, bacteria translocate into the bloodstream via the integument, respiratory system or gastrointestinal tract (1, 4, 5). Bacteria may have come from the immediate environment, or vertical transfer through the egg (6). Bacteria that survive in the blood may colonize the proximal growth plate of the rapidly growing femorae and tibiae, inducing necrosis leading to BCO-lameness (1, 5, 7). Stressors, or other factors that contribute to immunosuppression, can facilitate bacterial colonization and BCO spread in commercial poultry flocks (1, 8–13). In our research facility, *Staphylococcus agnetis* is the primary bacterium isolated from blood and infected bones of broilers when lameness is induced by growth on raised-wire-flooring (2, 5). Genomic analysis of *S. agnetis* isolates from chickens and dairy cattle demonstrate that the chicken isolates appear to be a clade arising from one branch of the cattle isolates (14). Multiple bacterial species have been historically identified from BCO lesions in different studies (1, 2, 11, 15–22) but there are few reports on characterization of genetic relatedness within isolates for a particular species obtained from chicken BCO lesions (20). Genomic analysis of a chicken isolate of *Staphylococcus aureus* suggested a genetic basis for the jump from human to chicken for an outbreak in the United Kingdom (23). Surveys of 20 broiler flock farms in Australia suggested that avian pathogenic *E. coli* was the main BCO isolate (24). Genetic analysis by multilocus sequence type (MLST), pulsed field gel electrophoresis and PCR phylogenetic grouping, of 15 *E. coli* isolates from 8 flocks in Brazil indicated significant diversity for vertebral osteomyelitis, and arthritis isolates, even within the same flock (25). The aim of this study was to characterize the genomes of BCO isolates from three different commercial broiler farms in Arkansas; to understand the origins of species causing BCO lameness in industry settings. Understanding the origins of the bacteria should influence strategies to reduce lameness outbreaks.

## Results and Discussion

### Diagnosis and Microbiological Sampling

In June of 2016 we surveyed two commercial broiler houses on separate farms experiencing outbreaks of BCO-lameness. Both houses had experienced a loss of cooling a week earlier, causing heat stress for several hours. The farms were in rural, central Arkansas, separated by 6.3 km, operated by the same integrator, and stocked from the same hatchery. The company veterinarian reported that samples from lame birds had been routinely submitted to a poultry health diagnostics laboratory and were primarily diagnosed as *E. coli.* For both farms we randomly collected lame birds for necropsy for BCO lesions. Blood and lesion swabs were collected from these birds, and house air was sampled, for bacterial species surveys. We used chromogenic diagnostic medium to assess whether multiple samples from the same bird appeared to yield significant numbers of a single colony type (size and color), indicating a single contributing species.

In Farm 1 the birds were 31 days old. We diagnosed and necropsied six lame birds (Table 1). KB1 and KB2 were symptomatic of spondylolisthesis/kinky-back (KB). KB1 had BCO lesions in T4, left tibia, and both femora. We obtained thousands (TNTC; too numerous to count) of small green colonies from the T4 sample that were determined to be *Enterococcus cecorum*. KB2 had BCO of only the left tibia, but no colonies were recovered from sampling from this site. We did recover approximately 50 green colonies from what appeared to be a normal T4 that were *E. cecorum*. Lame3 and Lame4 both had bilateral BCO of the femora and tibiae. Lame3 had TNTC white colonies from microbiological sampling of T4 that were *S. agnetis*. Lame5 had bilateral FHN, bilateral tibial dyschondroplasia (TD), and pericarditis. We recovered green colonies (20 from left and TNTC from right) from the TD lesions that were *E. cecorum*. Due to limited supplies there was no microbiological sampling for Lame4 and Lame6.

**Table 1.**
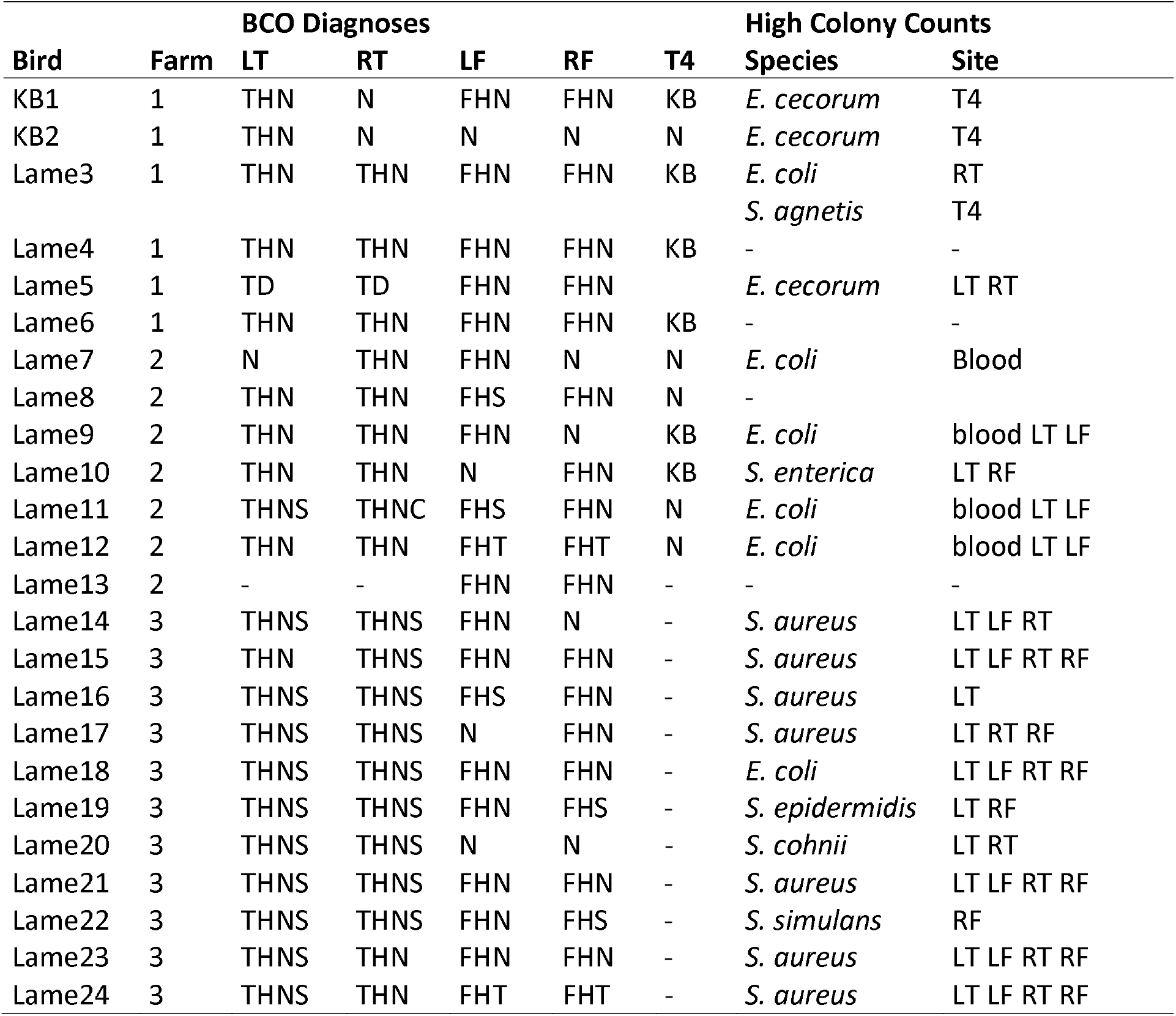
Microbiological sampling of bone and blood samples from two commercial broiler farms experiencing BCO outbreaks. BCO diagnoses are listed along with necropsy comments; RF-right femur, LF-left femur, RT-right tibia, LT-the left tibia, T4-vertebral joint, N-normal, THN-tibial head necrosis, THNS-THN severe, FHN-femoral head necrosis, KB-kinky back, TD-tibial dyschondroplasia, FHS-femoral head separation. High Colony Counts lists the Species diagnosis from 16S rRNA gene sequencing where a large number (>50) of same color colonies were recovered from one or more Site(s) indicating high probability of infection.

For Farm 2 the birds were 41 days of age. We diagnosed and necropsied seven lame birds (Table 1). Lame7 was diagnosed with BCO lesions of the left femur, and right tibia. We obtained 30 purple colonies that were *E. coli* from the blood sample. We diagnosed Lame8 with bilateral BCO of tibia and femur. No colonies were obtained from sampling this bird. Lame9 was diagnosed with BCO of bilateral tibia, and the left femur, with evident pericarditis. Approximately 100 purple colonies were produced from sampling of the left femur that were determined to be *E. coli.* Lame10 was diagnosed with BCO of both tibia and right femur. Lame10 also had pericarditis. We got 40 white colonies from the left tibia, and 70 white colonies from the right femur that were *Salmonella enterica*. Lame11 had BCO lesions on both tibiae and femora. We got TNTC purple colonies from the left tibia, three from the left femur, and 15 from blood that were determined to be *E. coli.* Lame12 was diagnosed with bilateral BCO of tibia and femur. We recovered approximately 500 purple colonies from blood, 40 purple colonies from the left tibia, and TNTC purple colonies from the left femur that were *E. coli.* Due to limited supplies there was no microbiological sampling for Lame13.

Average plate counts for air sampling were 80 and 125 for Farm 1 and Farm 2, respectively. The predominant species was *Staphylococcus cohnii* (~95%) with 3-4% *Staphylococcus lentus* and 1-2% *E. coli.* Air sampling during our broiler experiments at the University of Arkansas Poultry Research farm typically identify predominantly *S. cohnii* but not *E. coli* (26, 27).

In July 2019, we sampled a third commercial broiler farm (Farm 3) in Northwest Arkansas, more than 88 km from Farms 1 & 2 and operated by a different integrator supplied from a different hatchery. We sampled 11 lame birds at 35 days of age (Table 1). Lame14, was diagnosed with BCO of bilateral tibiae and right femur. We recovered numerous white colonies from the left femur, right tibia, and left tibia that were determined to be *Staphylococcus aureus*. Lame15 had BCO of all femorae and tibiae. Culture plates had numerous white colonies from all four sampled sites that were *S. aureus*. Lame16 had BCO of all tibiae and femorae. We recovered 10 green colonies from the right femur that were not analyzed, and 10 white colonies from the left tibia that were determined to be *S. aureus*. We diagnosed Lame17 with BCO of the right femur and both tibiae. We recovered numerous white colonies from all three sites that were *S. aureus*. Lame18 was diagnosed with bilateral BCO of the femorae and the tibiae. We got TNTC purple colonies from both femorae, and a few purple colonies from both tibiae, that were *E. coli.* Lame19 had BCO of all femorae and tibiae. Swabs gave only a few colonies of *S. epidermidis* that we assumed were contaminants during sampling. Lame20 had bilateral BCO of the tibiae. We recovered a few green colonies of *Staphylococcus cohnii* which were presumed contaminants during sampling. Lame21 had BCO of all femorae and tibiae. We isolated TNTC white colonies from all four sites that were *S. aureus*. Lame22 was diagnosed with bilateral BCO of femorae and tibiae. Microbial sampling only yielded only 10 green colonies from the right femur that were determined to be *Staphylococcus simulans*. Lame23 was diagnosed with bilateral BCO of femorae and tibiae. Culture plates had only white colonies, TNTC from both femora and tibiae, that were *S. aureus*. Lame24 had BCO of all femorae and tibiae. We recovered more than 100 white colonies from each of the four BCO lesions that were *S. aureus*.

### BCO Genome Assemblies

We chose to characterize genomes for four representative *E. coli* isolates: 1409 for Farm 1, 1413 from Farm 2, and 1512 and 1527 from one bird on Farm 3 (Table 2). A hybrid assembly for 1409 produced 5.05 Mbp in 23 contigs that organized into 4 DNA assembly graphs. We resolved the replicons using the long reads for contiguity analysis of the assembly graphs using the Bandage software. The resolved genome contains a 4.84 Mbp chromosome, with episomes of 113.6, 108.7, and 2.3. The predicted serotype was O16. The hybrid assembly for 1413 produced 5.37 Mbp in 59 contigs and 3 DNA assembly graphs. Unfortunately, the MinION reads were not of sufficient quality or length to complete a contiguity analysis of the entire genome, but did identify at least two episomes of 98.8 kbp and 2257 bp. The predicted serotype was O78. Draft assemblies were generated for *E. coli* 1512 and 1527. The assembly of 1512 contained 4.96 Mbp in 152 contigs with a N50 of 150 kbp. The assembly of 1527 was 4.90 Mbp in 179 contigs. The N50 was 97 Kbp with the largest contig of 258 Kbp. Both 1512 and 1527 were predicted to be serotype O78, like 1413. We generated draft assemblies for 14 *S. aureus* isolates from Farm 3 to examine genome diversity within a farm and within individual birds (Table 2). Two colonies from separate sample sites from seven lame birds were used for draft genome assembly (1510 & 1511, 1513 & 1514, 1515 & 1516, 1517 & 1518, 1519 & 1520, 1521 & 1522, 1523 & 1524). The assemblies (Table 2) ranged from 2.79 to 2.82 Mbp in 60 to 96 contigs (excluding contigs < 300 bp). The largest contigs were between 279 and 284 Kbp. N50 values ranged from 58 to 113 kbp. The L50 values ranged from 7 to 14 contigs. Each of the *S. aureus* assemblies had at least 3 circular contigs (episomes).

**Table 2.**
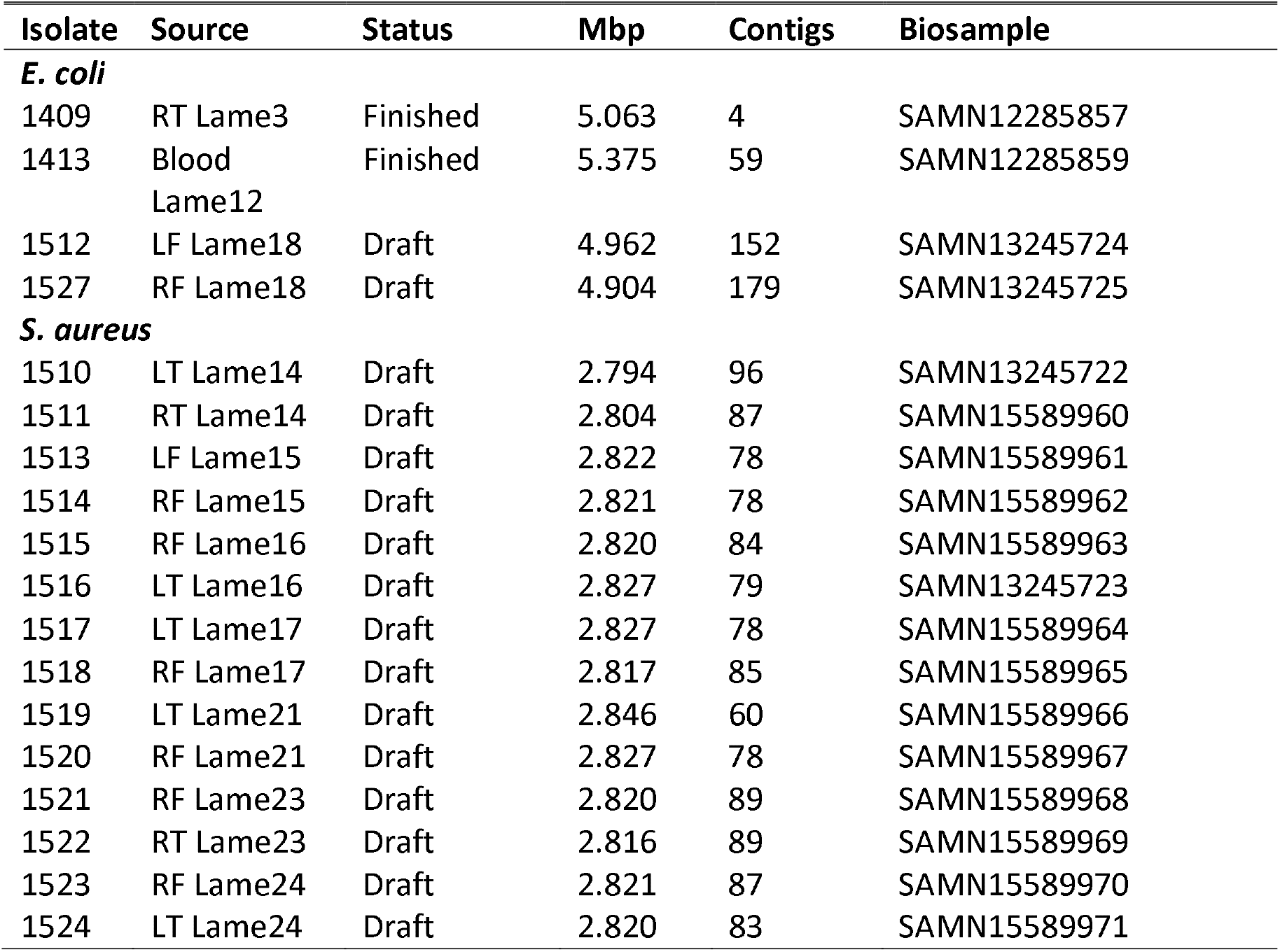
Bacterial genome assemblies produced in these analyses are listed by species, Isolate designation, host source, assembly genome Status and statistics (Mbp, Contig count), NCBI Biosample. Abbreviations are as in Table 1.

### Phylogenetic Comparison

To examine the phylogenetic relationships between *E. coli* isolates from the three farms, we identified the most closely related genomes, according to PATRIC, for 1409 (Farm 1), 1413 (Farm 2), 1512, and 1527 (Farm 3). We used the closest related genomes as surrogates, MOD1-EC6458 for 1409, PSUO78 for 1413, and ECO0667 for 1512/1527, to determine the placement of these isolates within the NCBI dendrogram for more than 18,000 *E. coli* genomes (https://www.ncbi.nlm.nih.gov/genome/167). This placed our isolates from the three farms in very distinct clades in the dendrogram for all E.coli genomes (data not shown). We downloaded 47 genomes representing the most closely related genomes identified by PATRIC for our four BCO *E. coli* genomes. We generated a phylogenetic tree for all 51 genomes using Average Nucleotide Identity (ANI), where genomes are identified by region and host/source (Figure 1 and Table S1). Isolate 1409 grouped with one chicken isolate from Pakistan and 4 isolates from chickens in China. The other closely related isolates in that same branch were from mammalian hosts from the USA, China, France or Mexico. ANI for this cluster is > 99.75%. Isolate 1413 clustered with 12 isolates from chickens, one from turkey meat, from the USA, United Kingdom, and Denmark, and three human isolates from Bolivia, Latvia and Mexico (ANI > 99.66%). Less-related genomes are from isolates from humans. Genomes from isolates 1512 and 1527 are virtually identical, with an ANI > 99.995%, which is not surprising since they were isolated from different anatomical sites in the same lame bird. Genomes for 1512 and 1527 clustered with those from chicken isolates from Poland, United Kingdom, and USA, along with genomes from isolates from humans in Japan, France and Estonia, pig isolates from China and USA, and a water sample from Arizona (ANI > 99.83%). Less-related genomes were from a colisepticemic turkey in Israel, Swiss chicken meat, a USA human isolate and deer feces from Pennsylvania. Therefore, while the clade for isolate 1413 seems to have a significant affinity for infecting poultry there are some human isolates. The clade for 1409 and the clade 1512/1527 both show a diversity of hosts including poultry and mammals. All three clades show a wide geographic distribution.

**Figure 1.**
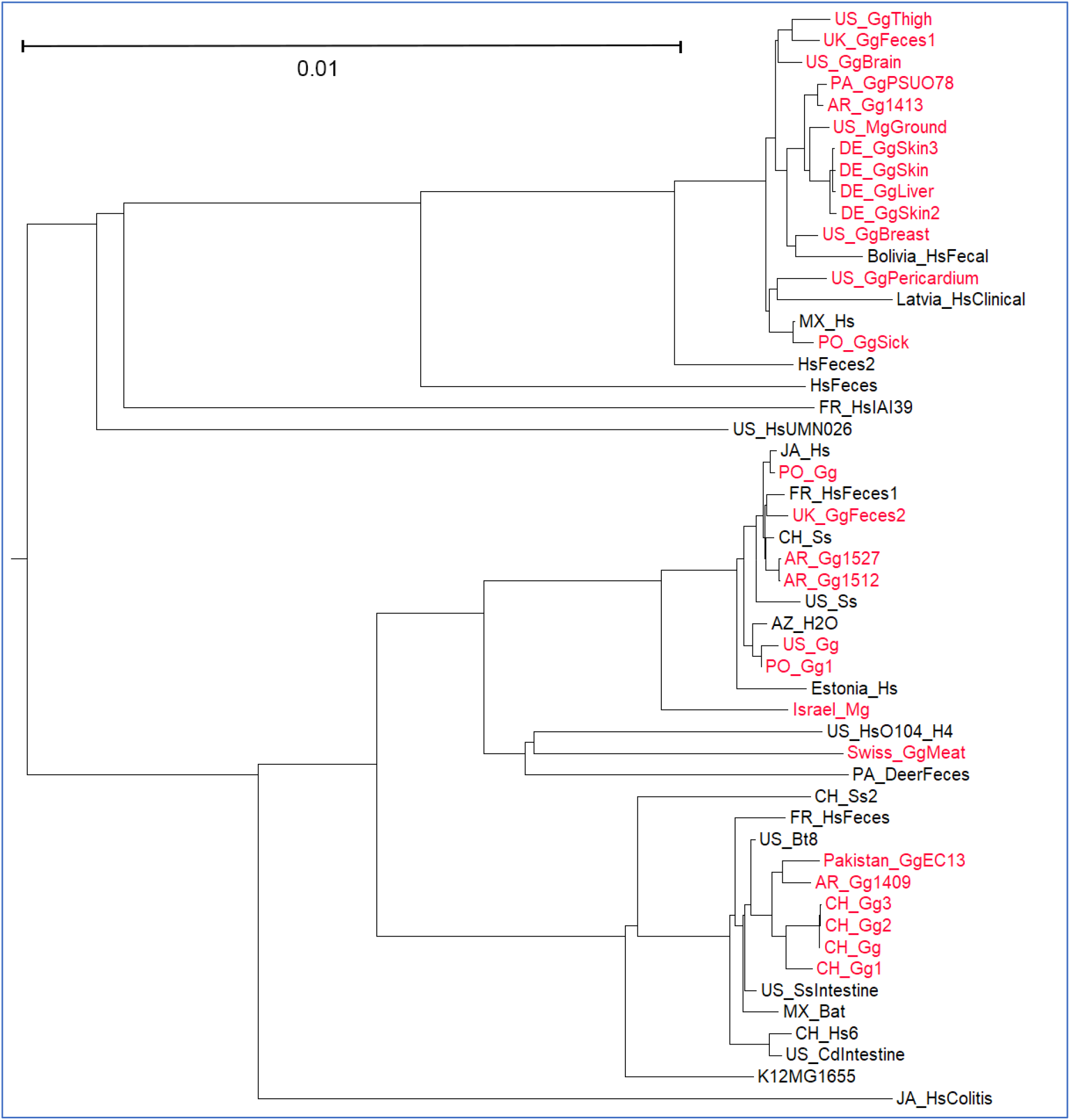
Phylogenetic tree for 51 *E. coli* genomes based on Average Nucleotide Identity. Key for isolate genomes is in Table S1. In brief, the prefix before the underline (_) indicates geographic location, first two characters after the underline indicate host, and remaining characters indicate source, strain, or isolate. Isolates in red are from poultry or poultry products.

The PATRIC closest-genome report for our new *S. aureus* chicken assemblies listed 11 *S. aureus* genomes for isolates obtained from chickens. Ten of these isolates cluster exclusively on one branch of a subtree in the NCBI genome dendrogram, and this cluster is next to a sister cluster of 13 human isolates (Figure 2). The close relationship of most of these chicken isolates had been previously demonstrated by Lowder *et al.* (23) using MLST to characterize *S. aureus* isolates from avian hosts. Those previous analyses suggested a specific host switch from humans to poultry and a close relationship among *S. aureus* isolated from the global chicken industry from 1970 to 2000. The genome of isolate ED98 was assembled as the type strain representing a BCO isolate from *S. aureus* from 1986 or 1987 in Ireland (23). We generated an ANI-based phylogenomic tree (Figure 3) for our 14 new *S. aureus* chicken BCO isolate genomes, the 11 chicken isolates from the ED98 clade, and included the two most closely related human isolates (based on PATRIC) from the sister clade in Figure 2. Our 14 BCO *S. aureus* genomes grouped together with an ANI > 99.998%, indicating a clonally-derived population. The 14 BCO *S. aureus* clustered with genomes for four isolates (B4-59C, B3-17D, B2-15A, B8-13D) from retail chicken meat from Tulsa, Oklahoma, in 2010 and a deep-wound lesion from a broiler from Poland in 2008 (ch23). Further distant are two additional Poland 2008 isolates (ch21 and ch22) from chicken wounds and lesions, and ED98 from the Ireland 1980s chicken BCO outbreak. Even further distant are ch9 from an infected chicken hock in 1999 in the USA, along with ch3 and ch5 which are recorded as chicken commensals from Belgium in 1976. The closest human isolates are CFBR-171 from a sputum sample from the USA in 2012 and GHA2 from a 2018 patient in Ghana. Additional details on these isolates are provided in Table S1. Our ANI based phylogenomics are in agreement with the work of Louder *et al.* (23), where they used MLST and mutational analysis of bi-allelic polymorphisms to demonstrate the tight relationship of 19 isolates of *S. aureus* from poultry in the USA, Japan, Denmark, Belgium, and United Kingdom. They had concluded that the “jump” to chickens likely was associated with the chicken commensals, ch3 and ch5, from Belgium. The 19 poultry isolates were derived from the human ST5 clade based and likely derived from human isolates circulating in Poland. The ED98 genome was assembled as representative of the chicken pathogens in 1986 or 1987. Limited genome comparisons prompted Lowder *et al.* (23) to conclude that the host switch by ED98 was associated with “acquisition of novel mobile genetic elements from an avian-specific accessory gene pool, and by the inactivation of several proteins important for human disease pathogenesis.” This was evidenced by their demonstration of enhanced resistance to killing by chicken heterophils.

**Figure 2.**
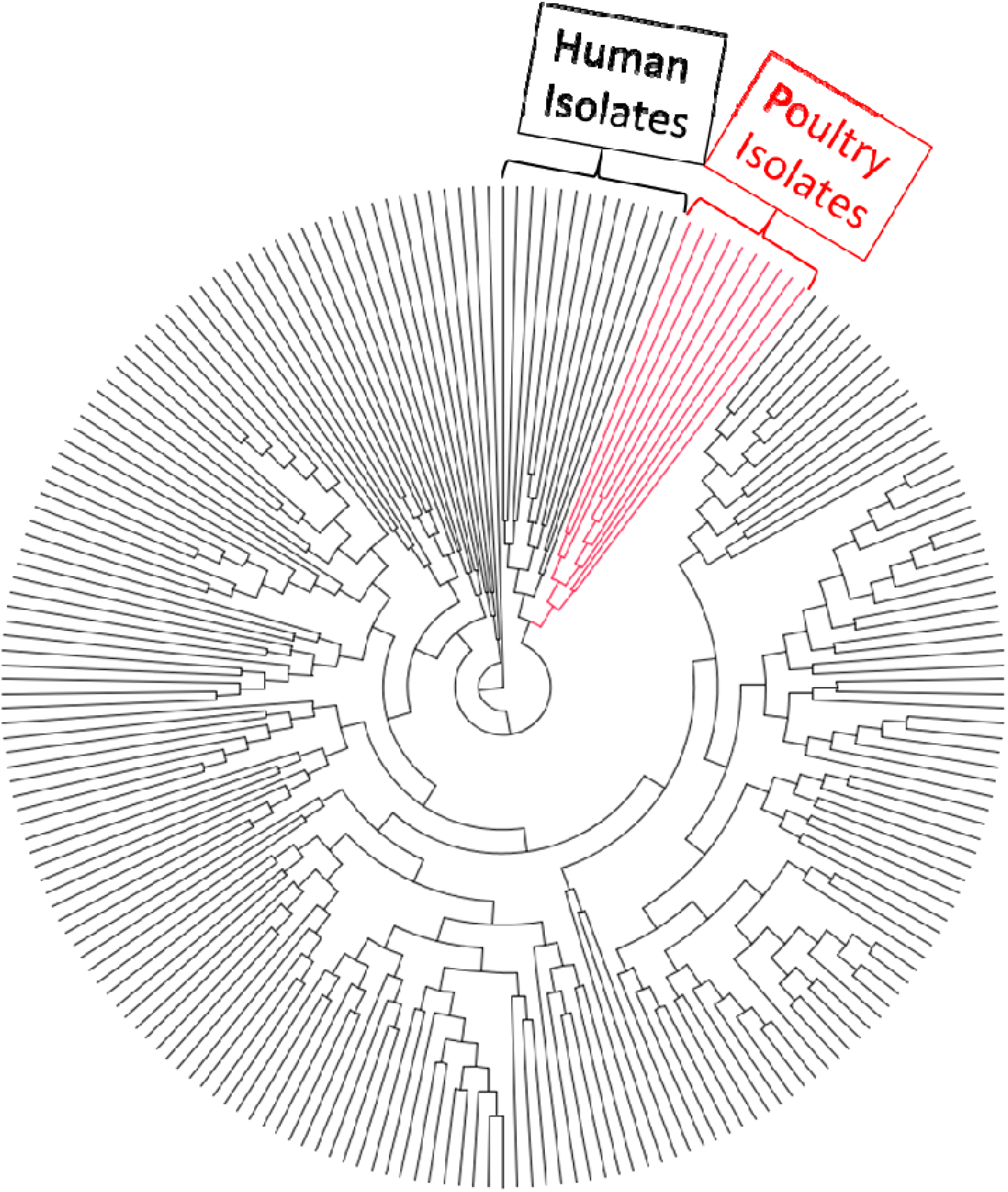
Phylogenetic subtree for 427 *S. aureus* genomes from NCBI based on whole genome BLAST comparisons. This subtree was an expanded view for the branch containing known poultry isolate genomes. The poultry isolate clade and a sister clade of human isolates are indicated.

**Figure 3.**
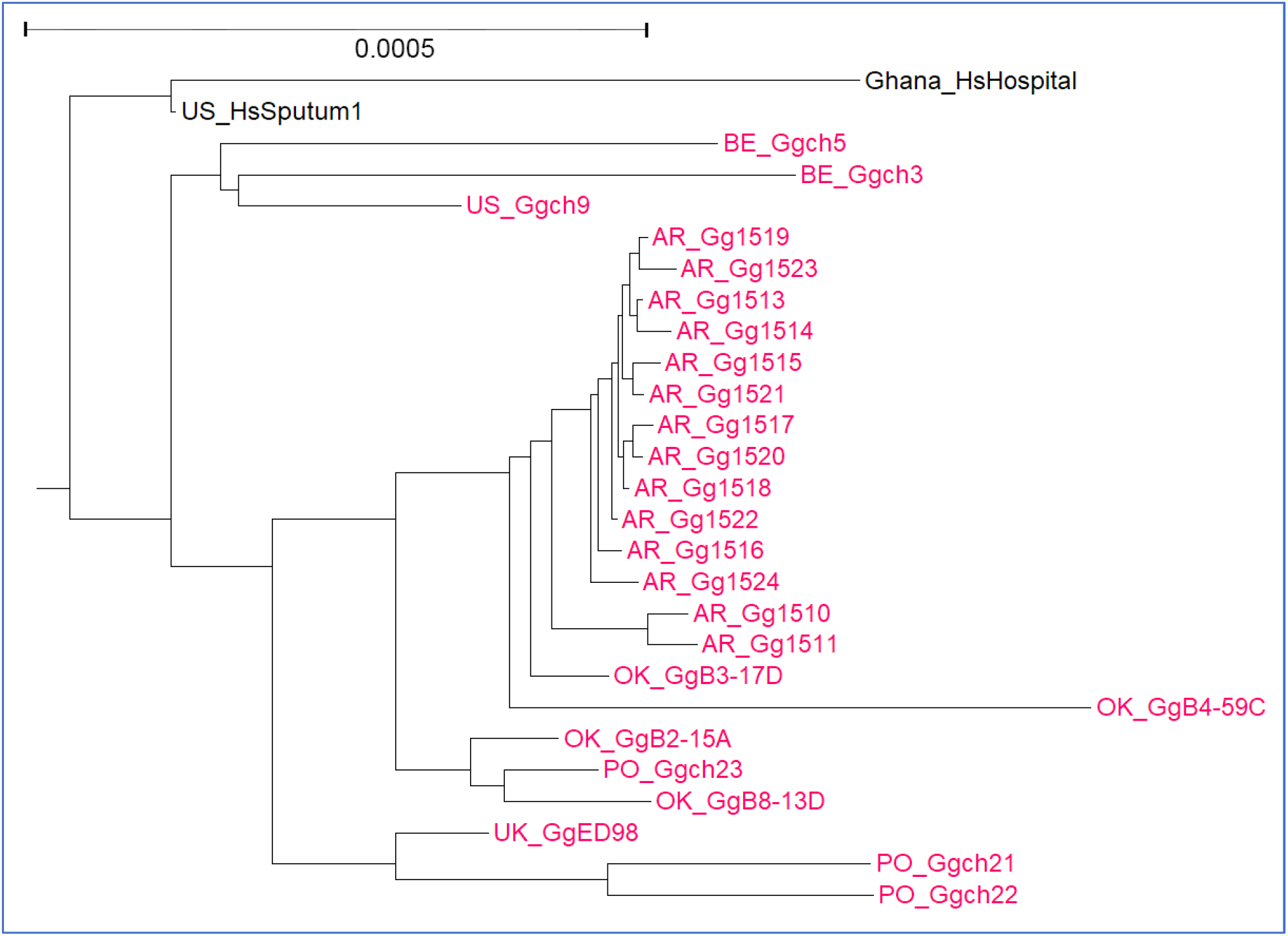
Phylogenetic tree for 27 *S. aureus* genomes based on Average Nucleotide Identity. Key for isolate genomes is in Table S1 and as described in legend to Figure 1. The tree compares 25 known isolates from poultry or poultry products (in red) with the closest related human isolates (in black).

In order to revisit the specific genomic changes in the clade of *S. aureus* isolates from chickens we used the RAST SEED Viewer proteome comparison tool to determine whether we could more precisely identify the genes distinguishing the human pathogens (GHA2 and CFBR-171), to either chicken commensal (ch3 and ch5), or chicken pathogens (ED98, B4-59C, and ch21). We used the ED98 predicted proteins as the reference, to identify polypeptides conserved over the entire polypeptide length at >80% identity in chicken pathogens, where the chicken commensals and human pathogens, encode a polypeptide of <80% identity. This filtering identified 33 protein encoding genes (PEG) from the main chromosome and none from the three plasmids in ED98 (Table 3 marked with *). Two clusters were identified associated with two mobile elements: a transposon and a *S. aureus* pathogenicity island (SaPI). There was also an additional short PEG (PEG 48). To further define any association of these 33 PEGs with the jump from humans to chickens we used tBLASTn to individually query the genomes of 11 *S. aureus* chicken isolates: 2 chicken commensals and 9 pathogens. We also performed a tBLASTn of 29 human *S. aureus* isolates including 10 from the sister clade in Figure 2 and an additional 19 representing clades flanking these two clades based on the NCBI genome tree. The tBLASTn included all 33 PEGs as well as immediate flanking or intervening PEGs (Table 3). The results show that PEG 48 is a hypothetical 56 residue polypeptide that appears only in the chicken pathogens. The transposon-related region (PEG 1641-1659) contained no PEG that was highly conserved in all the chicken pathogens while also less conserved in all the human isolates and the chicken commensals. Therefore PEG 48 or PEG 6141-1659 are not likely to be relevant to the jump from human to chicken pathogen. On the other hand, the SaPI region (PEG 755-773) contained a number of PEGs that appear highly conserved in all the chicken pathogens. Some of these PEGs (i.e., PEG 756-760, 767-771) are also highly conserved in some of the human pathogens, but lacking in the chicken commensals. The 4 hypothetical PEGs (774-778) downstream of the SaPI terminase all appear to be specific to the chicken pathogens. Close inspection of the results in Table 3 suggest that isolate ch9 appears to be intermediate between the chicken commensals, ch3 and ch5, and all of the chicken pathogens in the ED98 clade. This is concordant with our ANI phylogenomic analysis (Figure 3) where ch9 is basal relative to the other chicken pathogens. Thus, ch9 could derive from the earlier transitional state between the human pathogens to the chicken pathogens. We would anticipate that the genome evolved further from that earlier form as it adapted for colonization and pathogenesis in chickens. The data is consistent with the *S. aureus* genome evolving in two directions during the host switch from human to chickens: loss of the SaPI was associated with transition to a chicken commensal, and acquisition of new PEGs (perhaps 774-778) in the SaPI to become chicken-specialists. Louder *et al.* (23) had originally identified this SaPI region as associated with the jump to chickens but in their work they surveyed other avian isolates only by PCR amplification rather than our survey at the resolution of individual PEGs.

**Table 3.**
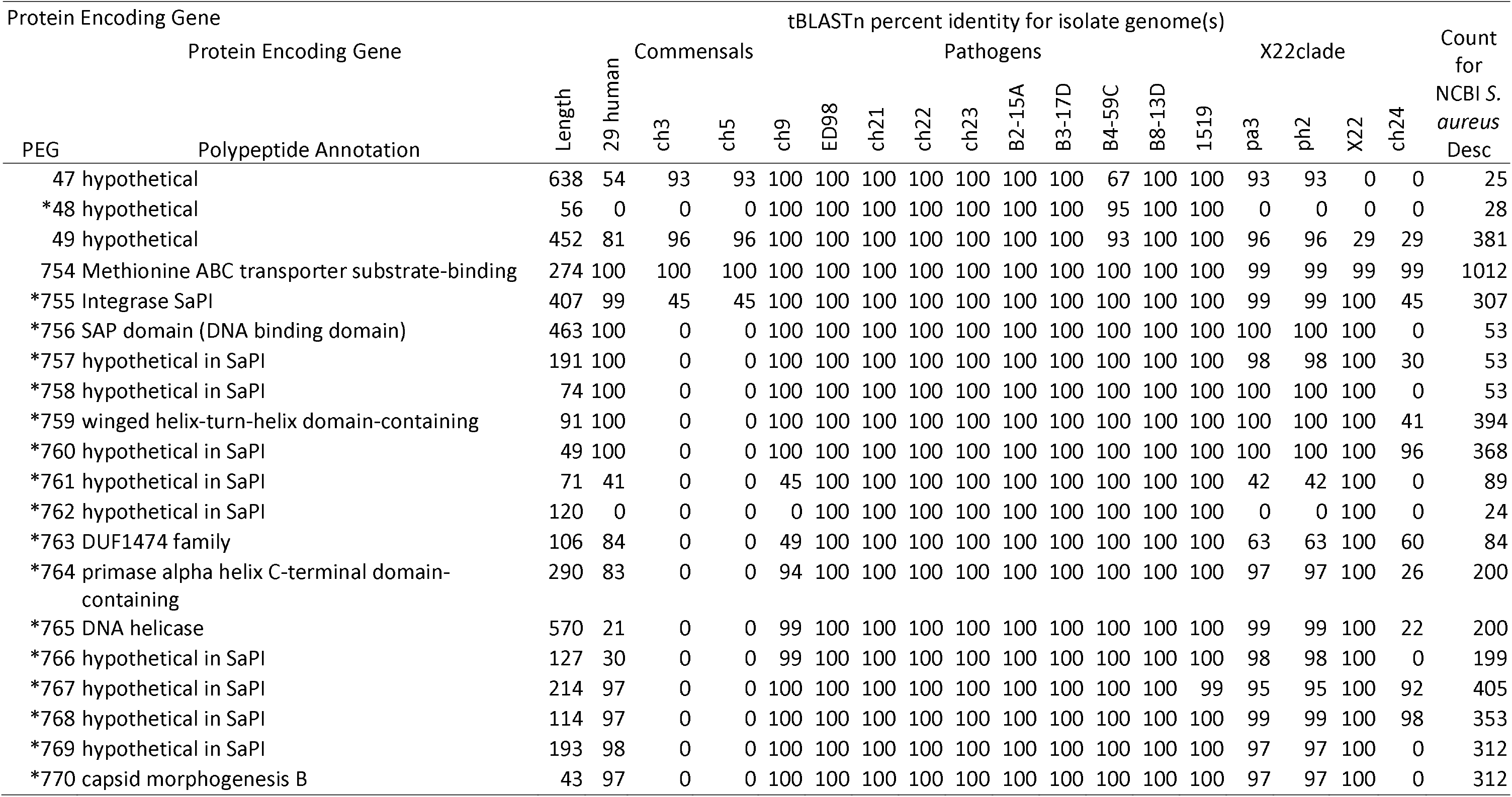
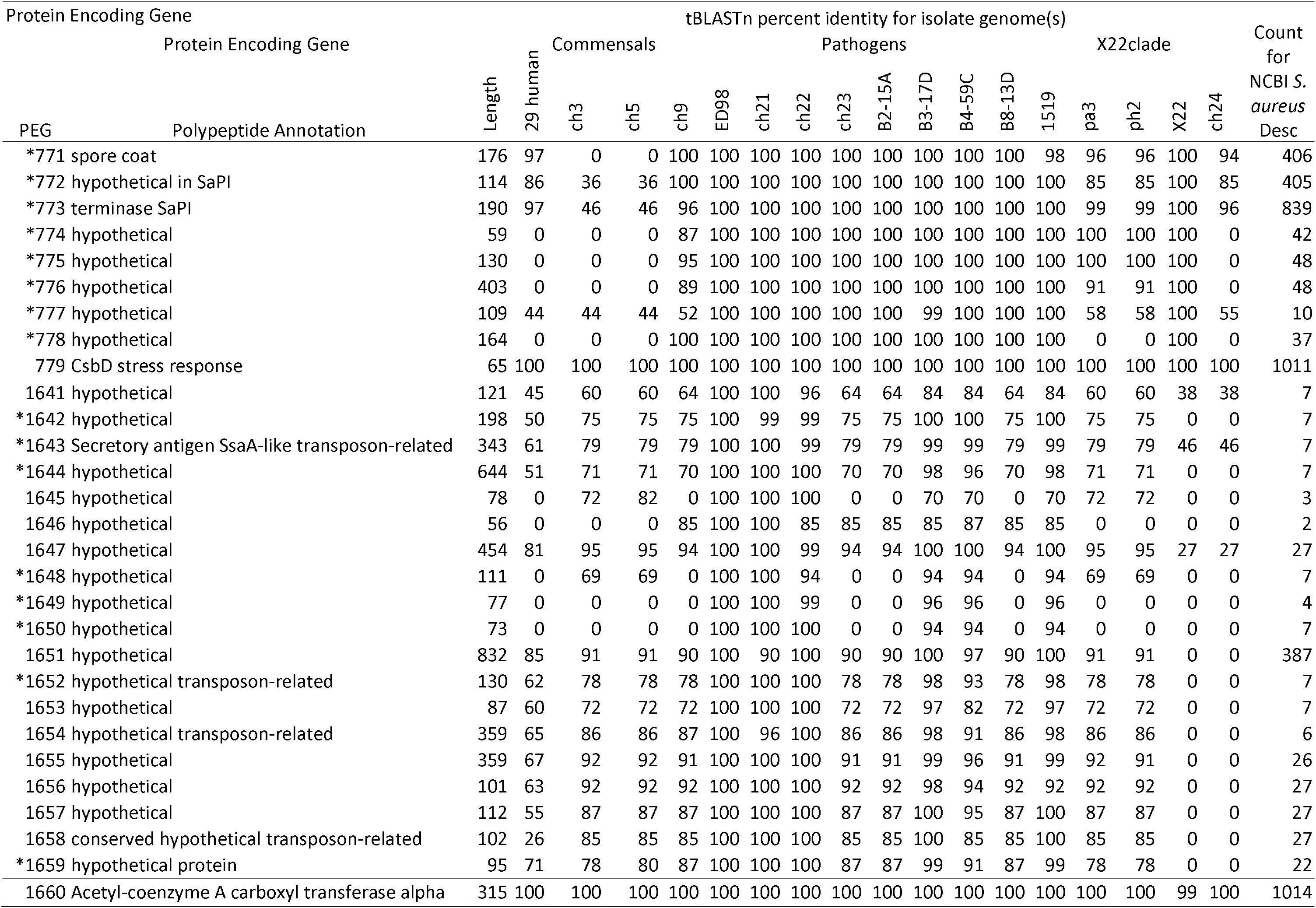
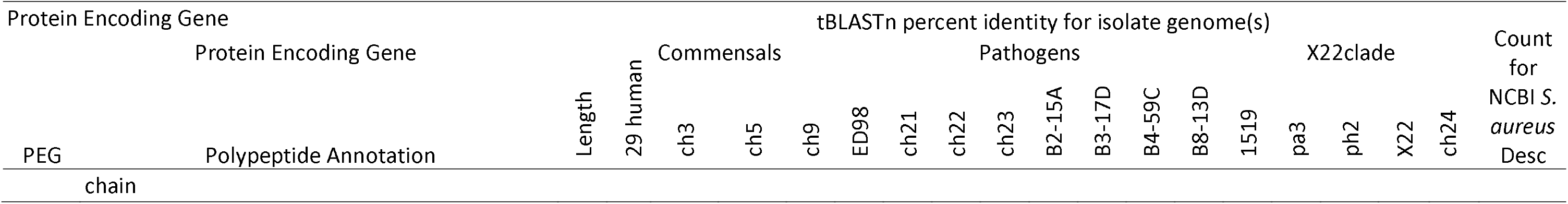
Percent identity from tBLASTn searches of *S. aureus* genomes using the polypeptide sequences for protein encoding genes from ED98. PEG originally identified as possibly chicken pathogen specific are indicated (*). Annotations are derived from SEED augmented by BLASTp queries at NCBI. The 29 human isolates are from the clades surrounding the ED98 clade as described in the text and identified in Table S1. Avian isolates include 2 chicken commensals (ch3 and ch5); 10 chicken pathogens from the ED98 clade; a partridge (pa3), pheasant (ph3) and unknown host (X22) from the X22 clade; and chicken isolate ch24 (not in the ED98 or X22 clade). Count for NCBI *S. aureus* Desc is the number of descriptions returned (max 5000) with >80% identity for an NCBI tBLASTn search of *S. aureus* (Taxonomy ID: 1280), which roughly approximates the number of genome entries encoding a highly related polypeptide. Additional genome details are in the text and Table S1.

SaPI are mobile elements that are packaged by “helper” virus assembly systems and integrate at a specific location (28, 29). The size of the SaPI element is determined by the packaging limits of the helper virus and SaPI accessory proteins. The SaPI in question is integrated between genes for a Methionine ABC transporter (PEG 754) and a CsbD stress response protein (PEG 779). We used tBLASTn of the *S. aureus* accessions in NCBI to survey for the number of entries with an integrase (PEG 755) homolog directly downstream of the Met ABC transporter homolog; with homolog threshold set at ≥80% identity for query polypeptide length. Out of 1012 genomes containing PEG 754 homologs (Table 3) there were 307 genomes with a PEG 755 homolog directly downstream; indicating a SaPI integration. Conversely, there were 568 genomes where a PEG 779 homolog directly followed PEG 754, consistent with no SaPI inserted in those genomes. Our tBLASTn searches also support that the SaPI mobile element extends from the integrase (PEG 755) through PEG 778, as we never found homologs of PEGs 777, 778, 780 or 781 directly downstream of PEG 754 (data not shown). Queries by tBLASTn using the PEG sequences from 754 to 779 demonstrate a high variability across the SaPI for whether the coding sequence is conserved in the genomes of other *S. aureus* isolates (see Count for NCBI *S. aureus* Desc column in Table 3). PEG 755, 759, 760, 764-773 appear to be present in many (Count range: 199-839) genomes, consistent with many having defined functions for SaPI mobilization and therefore conserved in many SaPI elemens. The 11 coding sequences for PEG 756-758, 761-763, and 774-778 are found in far fewer genomes (Count range: 10-89). We therefore used tBLASTn to identify those *S. aureus* accessions in NCBI with homologs to these 11 less-conserved PEGs. Only 8 accessions (ED98, ch21, ch22, B2-15A, B8-13D, B4-59C, B3-17D and X22) contained significant homologs (>80% identity over the entire query length) to all 11 coding sequences (Table 3). All are chicken pathogens from the same clade as ED98 with the exception of X22, which is a genome deposited by the China Animal Disease Control Center, but the host is not specified. Notably, the chicken commensals ch3 and ch5, have no SaPI downstream of the MetABC transporter (PEG 754). Therefore, this SaPI does not specify chicken colonization, but does correlate with pathogenicity in chickens. Clearly, more work needs to be focused on the actual functions of many of the genes in this SaPI and whether the chicken pathogenicity results from the combination of PEG 48 and the SaPI. Some of the PEGs in this SaPI have homologs in SaPIs in human isolates, however, the actual functions of the predicted polypeptides are not well understood (Table 3). One of the hypothetical PEGs, PEG 777, in this SaPI appears to be restricted to only chicken pathogens

We next explored whether there were signatures of selection in the evolution of the chicken pathogen genome progressing from ED98 to the present. We used the RAST SEED Viewer proteome comparison tool to analyze the evolution of this *S. aureus* chicken clade since 1987 (Table S2). We selected our assembly for 1519 as it was the largest assembly with the fewest contigs to represent the 2019 isolates from Farm 3. ED98 represents a 1986-1987 isolate in Ireland, ch21 is from Poland in 2008, and B4-59C is from 2010 in Tulsa, Oklahoma retail poultry meat. The SEED Viewer filter was set with the ED98 proteome as reference, to identify predicted proteins absent (<50% identity for query length) in one or more of the other three proteomes (Table S2). The analysis suggests that 32 proteins (31 phage and hypothetical proteins, and a efflux pump for Tetracycline resistance) were lost between 1987 and 2008. Eight phage and hypothetical proteins in ED98 and Ch21, were lost from the genome before appearance in Tulsa in 2010, and only 4 hypothetical proteins in ED98, ch21 and B4-59C, are absent in isolate 1519 in Northwest Arkansas in 2019. We then reversed the analysis with 1519 as the reference to identify new proteins that appeared in the lineage from ED98, through Ch21 to B4-59C to 1519. The analysis identified 35 polypeptide genes present in 1519 for which the other 3 genomes lack a polypeptide with 50% or greater overall identity. Twenty-eight are phage, hypothetical or plasmid-maintenance related. The remaining seven include a DUF1541 domain-containing polypeptide (PEG 32), a lead/cadmium/zinc/mercury/copper transporting ATPase (PEG 33), a partial coding sequence for phosphoglycerate kinase (PEG 421), and an aminoglycoside N6’-acetyltransferase (PEG 1919),. Two open reading frames (PEG 1916 and 1917) are not only new to the 1519 genome but have partially overlapping open reading frames. Thus, they may represent a frame shifted assembly error. However, reexamination of the templated alignment of the Illumina HiSeq reads with this particular 4512 bp contig showed deep coverage and no evidence for an assembly error. PEG 1916 annotates as a 51 residue SdrC, adhesin of unknown specificity. But BLASTp at NCBI annotates this as a partial (51 of 111 residues) sequence for an LPXTG cell wall anchor protein. PEG 1915 annotates as an 1125 residue, full length MSCRAMM SdrD homolog adhesin of unknown specificity. RAST annotates the 205 PEG 1917 as a methytransferase subunit for a Type I restriction system, but BLASTp searches at NCBI suggest an alternative as either an LPXT-cell wall anchor domain, and/or fibrinogen-binding protein. The assembly predicted this 4512 bp contig containing PEG 1915, 1916 and 1917 to be circular, so this may be a plasmid that encodes one or more adhesin functions. BLASTp with PEG 32, identified DUF1541 domain polypeptides of similar size in a wide range of different bacterial species. PEG33, the divalent cation transporter, is also found in many different *Staphylococcus species*. The aminoglycoside-N6’-acetyltransferase (gene 1919) has no significant BLASTp homologs in any *S. aureus* genome in NCBI, and the best homologs are 70% identical in isolates of *Staphylococcus sciuri, Staphylococcus lentus*, and *Staphylococcus fleurettii*. This gene is present in a 4357 bp contig that Unicycler could not circularize during assembly. However, the contig termini each contain portions of a plasmid recombination MobE mobilization protein that likely could be fused into one open reading frame using long read sequence data. The other genes in this contig are two hypothetical proteins, a tetracycline resistance predicted region, and an ArsR-family transcriptional regulator. However, this contig appears to possibly contain a mobile element affecting antibiotic resistance with the aminoglycoside transferase and the tetracycline resistance marker. Therefore, *S. aureus* 1519 appears to have obtained additional adhesins and possible antibiotic resistance markers since divergence from the *S. aureus* found in retail chicken meat in Oklahoma in 2010. Some of these appear to be associated with mobile elements.

Further evolution of this genome is evidenced by proteins highly conserved (>80%) in 1519 and B4-59C, but not (<50%) in ED98 and ch21. The Tulsa 2010 and Arkansas 2019 isolate genomes contain 16 proteins not found in ED98 and ch21; including a toxic shock syndrome toxin 1 (PEG 327), and a phage associated exotoxin superantigen (gene 329). Interestingly this region appears to be a probable SaPI. Inspection of neighboring genes identifies an integrase (PEG 335), terminase (PEG 326) and SaPI associated homologs (PEGs 325 and 330). PEG 327 and 329 are in a 97,219 bp contig predicted as circular that also encodes a number of genes for exotoxins and SaPI functions. The contig only contains two phage predicted proteins, so it may be a large plasmid containing many virulence determinants. Additional PEGs found in B4-59C and 1519 genomes but not ED98 and ch21 include a cluster (PEG 2179, 2180, and 2181) of homologs to hypothetical proteins found in SaPIs. However these PEGs are on a 3446 bp contig. Inspection of the SEED annotation of the B4-59C assembly identify flanking integrase and terminase homologs and other SaPI associated homologs. So, this region also may be a functional, mobile SaPI.

There were only 3 proteins identified in 1519, B4-59C and Ch21, but not in ED98; two are hypothetical (PEG 1197 and 1421) and the other a secretory antigen SsaA-like protein (PEG 49). This secretory antigen has been associated with transposons and also annotates as a CHAP domain protein, or putative cell wall lysis protein.

Our data demonstrate that this clade of *S. aureus* appears to be restricted to chickens for more than 40 years, and that it appears to have been in the Oklahoma/Arkansas region for more than a decade. Our genome comparisons of our recent isolates with the ED98 genome from 1986 shows that the genome has picked up additional virulence determinants (i.e., toxins), which appear to be associated with mobile SaPI. We do not know how this pathogen is transmitted to different farms or flocks. It could be vertically transmitted from hen to chicks. Alternatively, chicks could be exposed at the hatchery, or workers could spread the bacterium to farms through breakdowns in biosecurity.

In contrast the four *E. coli* genomes we characterized from three different farms show a different pattern. Isolates 1512 and 1527 are highly related as they came from different BCO lesions in the same lame bird. We have previously reported that, for individual lame birds, we recover the same species from multiple BCO lesions, and sometimes from the blood (2, 5). Our analyses of bacteria from BCO lesions in three farms is consistent with a predominant BCO pathogen within each farm, but multiple species may be contributing to BCO lameness within each facility. *E. coli* 1409, 1413, and 1512/1527 genomes are very distinct and come from very different clades. Reports from Brazil using virulence genes or MLST reported distinct *E. coli* genotypes within a flock (25). However, their data could not place the *E. coli* relative to those from non-chicken sources. The data for *E. coli* and BCO in chickens is different from the patterns for *S. aureus* where a single clade has been associated with chickens in Europe and the USA. In that respect we have similarly reported on a single clade of *S. agnetis* infecting chickens in the USA and Europe (14). The pattern we report from *E. coli* phylogenomics is most consistent with a generalist pathogen that easily jumps to different host species. Remarkably, two neighboring farms (Farm1 and Farm2) supplied by the same hatchery and operated by the same integrator, had very different *E. coli* (1409 and 1413) involved in BCO lameness outbreaks. This is more consistent with the *E. coli* on each farm originating from other hosts (zoonoses) or each farm could have “evolved” an *E. coli* BCO pathogen over many flocks and years.

## Conclusions

Overall, the *E. coli* isolates from BCO lesions in Arkansas appear to be highly diverse, as they derive from different clades that contain *E. coli* closely related to isolates from non-chicken hosts. Conversely, the *S. aureus* isolates appear to come from a clade of chicken-specific isolates associated exclusively with chicken hosts for at least four to five decades. Thus, the phylogenomics suggest that *E. coli* infecting chickens appears to be a generalist as highly related isolates are obtained from other hosts. S. aureus, which appears to be more of a specialist restricted to a single host. This distinction may derive from a difference in genome size as the *E. coli* genomes are roughly twice the size of the *S. aureus* genomes. The larger genome size would allow *E. coli* to retain a greater diversity of host-specificity virulence genes. Chicken pathogenicity of *S. aureus* appears to depend, in part, on a specific, mobile pathogenicity island in which the function of several genes are not defined. There is genomic evidence that the chicken pathogen clade continues to evolve, possibly driven by integration of additional different pathogenicity islands.

## Materials and Methods

### Microbiological Sampling and Bacterial Species Identification

Diagnosis of and sampling of BCO lesions and blood have been described (1, 2, 7, 30). Initial characterization of bacterial diversity by number of colonies of a particular color, was on CHROMagar Orientation (CO; DRG International, Springfield, NJ), and further refined by restreaking on CHROMagar Staphylococcus (CS; DRG International) (2, 5). Representative colonies were then diagnosed to species by 16S rRNA gene sequencing (2, 5).

Air sampling was by waving open CO plates within the building. CO and then further evaluated on CS plates. Representative colonies were typed to species as above.

#### Genomic DNA Isolation and Sequencing

Cultures were preserved in 40% glycerol at −80°C. Working stocks were maintained on tryptic soy agar slants at 4°C. For DNA extraction, staphylococci were grown in tryptic soy broth to log phase and DNA was isolated using as described (14). DNA isolation from *E. coli* used lysozyme treatment, followed by organic extractions (31). DNA was quantified using a GloMax® Multi Jr Detection System (Promega Biosystems Sunnyvale Inc., CA, USA) and purity evaluated with a Nanovue spectrophotometer (Healthcare Biosciences AB Uppsala, Sweden). DNA size was verified by agarose gel (1.5%) electrophoresis.

Library construction and Illumina MiSeq 2 × 250 sequencing were at the Michigan State University Genomics Core Facility. Libraries for Illumina HiSeqX 2 × 125 sequencing were prepared using a RipTide kit (iGenomX, Carlsbad, CA) and sequenced by Admera Health (South Plainfield, NJ). Long reads were generated using Oxford Nanopore-MinION bar-code kit, as described (14).

#### Genome Assembly and Analysis

De novo genome assemblies from short reads were generated as described (14). For hybrid assemblies the long reads were phase corrected using the Illumina short reads and Ratatosk v0.3 (https://github.com/DecodeGenetics/Ratatosk), before hybrid assembly with Unicycler v0.4.8 (32). Unicycler hybrid assembly graphs were further analyzed for contiguity in Bandage 0.8.1 (33) to discern and export replicons. The PATRIC (Pathosystems Resource Integration Center) webserver (34) was used for Unicycler assemblies, assembly annotation, and identification of similar genomes. Chromosome-level genomes were obtained from NCBI using genome_updater (https://github.com/pirovc/genome_updater). Average Nucleotide Identity (ANI) values were determined using pyANI 0.2.9 (35). ANI values were subtracted from 1 to generate distance matrices which were submitted to FastME 2.0 (36) using the BioNJ method to generate Newick trees. Archaeoptryx 0.9928 beta (37) was used to transform Newick trees into mid-point rooted graphic representations. Assemblies were annotated and compared using the Rapid Annotation using Subsystem Technologies (RAST) and SEED viewer (38, 39). Serotype prediction was using the ECTyper module at GalaxyTrakr.org. Local BLASTn and tBLASTn searches used BLAST 2.10.1+ (40).

## Acknowledgements

Support has been provided in part by grants from Cobb-Vantress, Inc., Zinpro LLC, and the Arkansas Biosciences Institute, the major research component of the Arkansas Tobacco Settlement Proceeds Act of 2000. BD was supported by KNOW (Leading National Research Centre), Scientific Consortium “Healthy Animal – Safe Food”, under of Ministry of Science and Higher Education No. 05-1/KNOW2/2015, and Department of Pathology and Veterinary Diagnostics, Institute of Veterinary Medicine, Warsaw University of Life Sciences–SGGW, Poland. The funders had no role in the design of this study, the interpretation of the results, or the contents of the manuscript. Thank you to Dr. Mark Hart and Dr. Karen Christensen for thoughtful comments regarding this manuscript.

