## Supplemental Tables 1 and 2 for "Analysis of Genomes of Bacterial Isolates from Lameness Outbreaks in Broilers"

Table S1. Genomes used for genomic analyses. Designation is the coding used in the phylogenomic trees, Country is the source of the isolate (if known; BE- Belgium, CH- China, DE- Denmark, FR- France, JA- Japan, MX- Mexico, PO- Poland, UK- United Kingdom), State is indicated for some USA isolates, Host is genus species and Source is the tissue/sampling from which the isolate was obtained. Assembly is the NCBI accession. BioProject is the NCBI registry for the isolate characterization.

| Designation | Strain | Host | Source | Country | State | Assembly | BioProject |
|---|---|---|---|---|---|---|---|
| *S. aureus* | | | | | | | |
| AR_Gg1510 | 1510 | *Gallus gallus* | Bone | USA | AR | JACEHY000000000 | PRJNA554887 |
| AR_Gg1511 | 1511 | *G. gallus* | Bone | USA | AR | JACEHW000000000 | PRJNA554887 |
| AR_Gg1513 | 1513 | *G. gallus* | Bone | USA | AR | JACEHV000000000 | PRJNA554887 |
| AR_Gg1514 | 1514 | *G. gallus* | Bone | USA | AR | JACEHU000000000 | PRJNA554887 |
| AR_Gg1515 | 1515 | *G. gallus* | Bone | USA | AR | JACEHT000000000 | PRJNA554887 |
| AR_Gg1516 | 1516 | *G. gallus* | Bone | USA | AR | JACEHX000000000 | PRJNA554887 |
| AR_Gg1517 | 1517 | *G. gallus* | Bone | USA | AR | JACEHS000000000 | PRJNA554887 |
| AR_Gg1518 | 1518 | *G. gallus* | Bone | USA | AR | JACEHR000000000 | PRJNA554887 |
| AR_Gg1519 | 1519 | *G. gallus* | Bone | USA | AR | JACEHQ000000000 | PRJNA554887 |
| AR_Gg1520 | 1520 | *G. gallus* | Bone | USA | AR | JACEHP000000000 | PRJNA554887 |
| AR_Gg1521 | 1521 | *G. gallus* | Bone | USA | AR | JACEHO000000000 | PRJNA554887 |
| AR_Gg1522 | 1522 | *G. gallus* | Bone | USA | AR | JACEHN000000000 | PRJNA554887 |
| AR_Gg1523 | 1523 | *G. gallus* | Bone | USA | AR | JACEHM000000000 | PRJNA554887 |
| AR_Gg1524 | 1524 | *G. gallus* | Bone | USA | AR | JACEHL000000000 | PRJNA554887 |
| OK_GgB2-15A | B2-15A | *G. gallus* | Retail meat | USA | OK | GCA_007726495.1 | PRJNA555718 |
| OK_GgB3-17D | B3-17D | *G. gallus* | Retail meat | USA | OK | GCA_007726565.1 | PRJNA555718 |
| OK_GgB4-59C | B4-59C | *G. gallus* | Retail meat | USA | OK | GCA_007726525.1 | PRJNA555718 |
| OK_GgB8-13D | B8-13D | *G. gallus* | Retail meat | USA | OK | GCA_007726545.1 | PRJNA555718 |
| PO_Ggch21 | ch21 | *G. gallus* | Deep wound | Poland | | GCA_003343155.1 | PRJNA344860 |
| PO_Ggch22 | ch22 | *G. gallus* | Deep wound | Poland | | GCA_003350605.1 | PRJNA344860 |
| PO_Ggch23 | ch23 | *G. gallus* | Deep wound | Poland | | GCA_003336545.1 | PRJNA344860 |
| BE_Ggch3 | ch3 | *G. gallus* | Commensal | Belgium | | GCA_003336625.1 | PRJNA344860 |
| BE_Ggch5 | ch5 | *G. gallus* | Commensal | Belgium | | GCA_003336635.1 | PRJNA344860 |
| US_Ggch9 | ch9 | *G. gallus* | Hock | USA | | GCA_003336495.1 | PRJNA344860 |
| UK_GgED98 | ED98 | *G. gallus* | Bone | Ireland | | GCA_000024585.1 | PRJNA39547 |
| | X22 | *Unknown* | Unknown | China | | GCA_007998145.1 | PRJNA558859 |
| | ph2 | Pheasant | Tenosynovitis | Scotland | | GCA_003336565.1 | PRJNA344860 |
| | pa3 | Partridge | Liver | Scotland | | GCA_003336575.1 | PRJNA344860 |

| Designation | Strain | Host | Source | Country | State | Assembly | BioProject |
|---|---|---|---|---|---|---|---|
| Ghana_HsHospital | GHA2* | *Homo sapiens* | Hospital | Ghana | | GCA_008630855.1 | PRJNA564764 |
| US_HsSputum | CFBR-122* | *H. sapiens* | Sputum | USA | | GCA_003720355.1 | PRJNA480016 |
| | 3688STDY6124906* | *H. sapiens* | Unknown | Thailand | | GCA_900126025.1 | PRJEB9575 |
| | 3688STDY6125000* | *H. sapiens* | Unknown | Thailand | | GCA_900124915.1 | PRJEB9575 |
| | BCH-SA-12* | *H. sapiens* | Throat | USA | | GCA_003720885.1 | PRJNA480016 |
| | CFBR-102* | *H. sapiens* | Sputum | USA | | GCA_003720115.1 | PRJNA480016 |
| | CFBR-149* | *H. sapiens* | Sputum | USA | | GCA_003720135.1 | PRJNA480016 |
| | CFBR-171* | *H. sapiens* | Sputum | USA | | GCA_003719905.1 | PRJNA480016 |
| | CFSA134* | *H. sapiens* | Sputum | USA | | GCA_002123885.1 | PRJNA380429 |
| | DAR1813* | *H. sapiens* | Blood | USA | | GCA_000609765.1 | PRJNA228330 |
| | DAR1890* | *H. sapiens* | Fluid | USA | | GCA_000609945.1 | PRJNA228339 |
| | DAR3156* | *H. sapiens* | Bone | Argentina | | GCA_000610525.1 | PRJNA228372 |
| | DAR3157* | *H. sapiens* | Sepsis | Argentina | | GCA_000610545.1 | PRJNA228373 |
| | DAR3166* | *H. sapiens* | Bone | Argentina | | GCA_000610605.1 | PRJNA228376 |
| | DAR3178* | *H. sapiens* | Bone | Argentina | | GCA_000610685.1 | PRJNA228381 |
| | F29982* | *H. sapiens* | Nares | USA | | GCA_000571715.1 | PRJNA225050 |
| | F41882* | *H. sapiens* | Nares | USA | | GCA_000559345.1 | PRJNA224323 |
| | FDAARGOS_14* | *H. sapiens* | Unknown | USA | NY | GCA_001018775.2 | PRJNA231221 |
| | FDAARGOS_16* | *H. sapiens* | Unknown | USA | NY | GCA_001019355.2 | PRJNA231221 |
| | FDAARGOS_359* | *H. sapiens* | Wound | USA | MD | GCA_002554295.1 | PRJNA231221 |
| | H27777* | *H. sapiens* | Nares | USA | | GCA_000561365.1 | PRJNA224442 |
| | M0455* | *H. sapiens* | ICU | USA | | GCA_000361205.1 | PRJNA173479 |
| | M6K089* | *H. sapiens* | Skin | Japan | | GCA_003421965.1 | PRJDB5246 |
| | M6K136* | *H. sapiens* | Skin | Japan | | GCA_003422345.1 | PRJDB5246 |
| | NCTC10804* | unknown | Unknown | UK | | GCA_900458375.1 | PRJEB6403 |
| | NCTC9393* | *H. sapiens* | Unknown | UK | | GCA_900458005.1 | PRJEB6403 |
| | RN6607* | *H. sapiens* | Newborn nursery | USA | NY | GCA_000597965.1 | PRJNA240093 |
| | SAM-7* | *H. sapiens* | Throat | Lebanon | | GCA_003038745.1 | PRJNA437720 |
| | W76127* | *H. sapiens* | Nares | USA | | GCA_000562505.1 | PRJNA224506 |
| *E. coli* | | | | | | | |
| AR_Gg1409 | 1409 | *G. gallus* | Bone | USA | AR | JACGTG000000000 | PRJNA554886 |
| AR_Gg1413 | 1413 | *G. gallus* | Bone | USA | AR | JACGTF000000000 | PRJNA554886 |
| AR_Gg1512 | 1512 | *G. gallus* | Bone | USA | AR | JACGTE000000000 | PRJNA554886 |

| Designation | Strain | Host | Source | Country | State | Assembly | BioProject |
|---|---|---|---|---|---|---|---|
| AR_Gg1527 | 1527 | *G. gallus* | Bone | USA | AR | JACGTD000000000 | PRJNA554886 |
| Bolivia_HsFecal | 286A | *H. sapiens* | Feces | Bolivia | | GCA_003850735.1 | PRJNA427943 |
| CH_Gg | 12c7 | *G. gallus* | | China | | GCA_003009015.1 | PRJNA417344 |
| CH_Gg1 | YH17134 | *G. gallus* | | China | | GCA_002959165.1 | PRJNA434044 |
| CH_Gg2 | 12c8 | *G. gallus* | | China | | GCA_003008775.1 | PRJNA417344 |
| CH_Gg3 | 12c5 | *G. gallus* | | China | | GCA_003009715.1 | PRJNA417344 |
| CH_Hs6 | A61 | *H. sapiens* | Feces | China | | GCA_003302635.1 | PRJNA400107 |
| CH_Ss | E565 | *Sus scrofa* | | China | | GCA_003328175.1 | PRJNA450836 |
| CH_Ss2 | SEC470 | *S. scrofa* | Feces | China | | GCA_000987875.1 | PRJNA244370 |
| DE_GgLiver | E44 | *G. gallus* | Liver | Denmark | | GCA_001652345.1 | PRJNA321591 |
| DE_GgSkin | L7S7 | *G. gallus* | Skin | Denmark | | GCA_003015065.1 | PRJNA438734 |
| DE_GgSkin2 | L3S3 | *G. gallus* | Skin | Denmark | | GCA_003013555.1 | PRJNA438662 |
| DE_GgSkin3 | L5S5 | *G. gallus* | Skin | Denmark | | GCA_003015075.1 | PRJNA438735 |
| Estonia_Hs | EEIVKB55 | *H. sapiens* | Clinical sample | Estonia | | GCA_006238365.1 | PRJNA528606 |
| FR_HsFeces | CEREMI_E32 | *H. sapiens* | Feces | France | | GCA_900536595.1 | PRJEB28341 |
| FR_HsFeces1 | 884A | *H. sapiens* | Feces | France | | GCA_900499885.1 | PRJEB28020 |
| FR_HsIAI39 | IAI39 | *H. sapiens* | | France | | GCA_000026345.1 | PRJNA33411 |
| HsFeces | 2012C-4502 | *H. sapiens* | Feces | | | GCA_003018255.1 | PRJNA218110 |
| HsFeces2 | 2014C-3338 | *H. sapiens* | | | | GCA_003018095.1 | PRJNA218110 |
| Israel_Mg | 789 | *Meleagris gallopavo* | Blood | Israel | | GCA_000819645.1 | PRJNA262513 |
| JA_Hs | SMEc189 | *H. sapiens* | | Japan | | GCA_006535915.1 | PRJDB8148 |
| JA_HsColitis | SAKAI (EHEC) | *H. sapiens* | Hemorrhagic colitis | Japan | | GCA_000008875.1 | PRJNA226 |
| K12MG1655 | MG1655 | Unknown | | | | GCA_000005845.2 | PRJNA603343 |
| Latvia_HsClinical | LVSTRB103 | *H. sapiens* | Clinical sample | Latvia | | GCA_006236595.1 | PRJNA528606 |
| MX_Bat | MOD1-EC908 | *Tadarida brasiliensis* | Feces | Mexico | | GCA_002456375.1 | PRJNA230969 |
| MX_Hs | MOD1-EC6621 | *H. sapiens* | Feces | Mexico | | GCA_002485345.1 | PRJNA230969 |
| PA_DeerFeces | PSUO103 | *Odocoileus virginianus* | Feces | USA | PA | GCA_002215155.1 | PRJNA314794 |
| PA_GgPSUO78 | PSUO78 | *G. gallus* | Peritoneum | USA | | GCA_002215115.1 | PRJNA287566 |
| Pakistan_GgEC13 | EC_13 | *G. gallus* | Infection | Pakistan | | GCA_004284075.1 | PRJNA522294 |

| Designation | Strain | Host | Source | Country | State | Assembly | BioProject |
|---|---|---|---|---|---|---|---|
| PO_Gg | 019PP2015 | *G. gallus* | | Poland | | GCA_001709145.1 | PRJNA319144 |
| PO_GgSick | 012PP2015 | *G. gallus* | Sick bird | Poland | | GCA_001696335.1 | PRJNA319144 |
| PO_Gg2 | 022PP2016 | *G. gallus* | | Poland | | GCA_001758245.1 | PRJNA319144 |
| Swiss_GgMeat | S51 | *G. gallus* | Meat | Switzerland | | GCA_001660565.1 | PRJNA323827 |
| UK_GgFeces1 | VREC0540 | *G. gallus* | Feces | United Kingdom | | GCA_900490165.1 | PRJEB8774 |
| UK_GgFeces2 | VREC0637 | *G. gallus* | Feces | United Kingdom | | GCA_900482085.1 | PRJEB8774 |
| US_Bt8 | KCJK8229 | *B. taurus* | Feces | USA | | GCA_004792865.1 | PRJNA420036 |
| US_CdIntestine | MOD1-EC5097 | *Canis domesticus* | Intestine | USA | NY | GCA_002232435.1 | PRJNA230969 |
| US_Gg | MOD1-EC6339 | *G. gallus* | Egg | USA | AL | GCA_002512585.1 | PRJNA230969 |
| US_GgBrain | MOD1-EC6094 | *G. gallus* | Brain | USA | | GCA_002537555.1 | PRJNA230969 |
| US_GgBreast | CVM N17EC0744 | *G. gallus* | Breast meat | USA | | GCA_003793955.1 | PRJNA292663 |
| US_GgPericardium | MOD1-EC5115 | *G. gallus* | Pericardial Sac | USA | PA | GCA_002231405.1 | PRJNA230969 |
| US_GgThigh | CVM N17EC0412 | *G. gallus* | Chicken Thighs | USA | | GCA_003794735.1 | PRJNA292663 |
| US_HsFeces | 2011C-3493 | *H. sapiens* | Feces | USA | | GCA_000299455.1 | PRJNA81095 |
| US_Hs | UMN026 | *H. sapiens* | | USA | | GCA_000026325.2 | PRJNA33415 |
| US_MgGround | CVM N17EC1100 | *M. gallopavo* | Ground meat | USA | | GCA_003774815.1 | PRJNA292663 |
| US_Ss | MOD1-EC5757 | *S. scrofa* | Ileum | USA | SD | GCA_002474525.1 | PRJNA230969 |
| US_SsIntestine | MOD1-EC6458 | *S. scrofa* | Jejunum | USA | | GCA_002464015.1 | PRJNA230969 |
| AZ_H2O | MOD1-EC5915 | | Water | USA | AZ | GCA_002534895.1 | PRJNA230969 |

*- 29 *S. aureus* isolates used in tBLASTn analyses in Table 3.

Table S2. Evolution of proteomes in four genomes of *S. aureus* infecting chickens. The RAST SEED viewer was used to identify genes present in *S.aureus* 1519 where the predicted polypeptide had a % identity less than 50% in one or more of the genomes for the indicated isolates. PEG is the RAST 1519 annotation gene number, Length is for the predicted polypeptide in 1519, and function is the annotation from RAST and/or NCBI.

| % Identity | | | 1519 | | |
|---|---|---|---|---|---|
| ED98 | Ch21 | B4-59C | PEG | Length | function |
| 100.0 | 100.0 | 0.0 | 6 | 148 | hypothetical protein |
| 100.0 | 100.0 | 0.0 | 7 | 83 | hypothetical protein |
| 100.0 | 100.0 | 0.0 | 16 | 112 | hypothetical protein |
| 0.0 | 0.0 | 0.0 | 32 | 182 | DUF1541 domain-containing protein |
| 36.4 | 35.9 | 36.4 | 33 | 688 | Lead, cadmium, zinc and mercury transporting ATPase; Copper-translocating P-type ATPase |
| 0.0 | 0.0 | 0.0 | 117 | 43 | hypothetical protein |
| 0.0 | 0.0 | 0.0 | 153 | 38 | hypothetical protein |
| 0.0 | 0.0 | 100.0 | 296 | 97 | hypothetical protein |
| 0.0 | 34.4 | 100.0 | 327 | 235 | Toxic shock syndrome toxin 1 (TSST-1) |
| 0.0 | 0.0 | 100.0 | 328 | 55 | hypothetical protein |
| 0.0 | 55.0 | 100.0 | 329 | 41 | Exotoxin, phage associated, enterotoxin SEB (partial) |
| 0.0 | 0.0 | 100.0 | 331 | 140 | Phage protein |
| 0.0 | 0.0 | 100.0 | 332 | 390 | Phage protein |
| 0.0 | 0.0 | 0.0 | 421 | 40 | Phosphoglycerate kinase (partial) |
| 45.9 | 45.9 | 100.0 | 500 | 48 | hypothetical protein |
| 0.0 | 100.0 | 100.0 | 562 | 49 | Secretory antigen SsaA-like protein transposon-related |
| 0.0 | 0.0 | 0.0 | 595 | 44 | hypothetical protein |
| 36.0 | 36.0 | 100.0 | 729 | 222 | hypothetical protein |
| 26.1 | 26.1 | 100.0 | 732 | 242 | hypothetical protein |
| 0.0 | 0.0 | 100.0 | 733 | 71 | Phage protein |
| 100.0 | 100.0 | 0.0 | 1021 | 128 | hypothetical protein |
| 0.0 | 100.0 | 100.0 | 1197 | 45 | hypothetical protein |
| 100.0 | 100.0 | 0.0 | 1210 | 39 | hypothetical protein |
| 0.0 | 0.0 | 0.0 | 1289 | 64 | hypothetical protein |
| 0.0 | 100.0 | 100.0 | 1421 | 44 | hypothetical protein |
| 100.0 | 98.6 | 0.0 | 1553 | 74 | Hypothetical protein, phi-ETA orf24 homolog |
| 0.0 | 0.0 | 0.0 | 1554 | 72 | Phage protein |
| 37.4 | 36.1 | 37.4 | 1555 | 414 | Phage DNA helicase |

| % Identity | | | 1519 | | |
|---|---|---|---|---|---|
| ED98 | Ch21 | B4-59C | PEG | Length | function |
| 0.0 | 0.0 | 0.0 | 1556 | 119 | Phage protein |
| 0.0 | 0.0 | 0.0 | 1557 | 255 | Phage replication initiation protein |
| 95.9 | 97.7 | 0.0 | 1558 | 225 | Hypothetical protein, PV83 orf19 homolog |
| 100.0 | 99.5 | 0.0 | 1560 | 213 | Phage-associated recombinase |
| 100.0 | 100.0 | 0.0 | 1561 | 160 | ORF027 |
| 98.8 | 97.7 | 39.5 | 1563 | 87 | Hypothetical protein, PVL orf39 homolog |
| 97.3 | 98.6 | 0.0 | 1565 | 74 | Hypothetical protein, PV83 orf12 homolog |
| 98.2 | 98.2 | 0.0 | 1566 | 57 | Uncharacterized protein pCM2_0059 |
| 100.0 | 100.0 | 0.0 | 1567 | 97 | phage protein |
| 100.0 | 81.3 | 0.0 | 1568 | 251 | Phage antirepressor protein |
| 100.0 | 100.0 | 0.0 | 1569 | 62 | hypothetical protein |
| 100.0 | 100.0 | 0.0 | 1570 | 201 | Phage protein |
| 0.0 | 0.0 | 0.0 | 1572 | 65 | Phage protein |
| 0.0 | 0.0 | 0.0 | 1573 | 257 | Phage antirepressor protein |
| 0.0 | 0.0 | 0.0 | 1574 | 79 | hypothetical protein |
| 100.0 | 93.4 | 0.0 | 1576 | 110 | Phage protein |
| 99.7 | 100.0 | 0.0 | 1577 | 553 | DNA adenine methylase |
| 100.0 | 100.0 | 25.7 | 1578 | 350 | Phage integrase |
| 0.0 | 0.0 | 0.0 | 1719 | 41 | hypothetical protein |
| 0.0 | 0.0 | 100.0 | 1760 | 52 | hypothetical protein transposon-related |
| 0.0 | 0.0 | 0.0 | 1862 | 158 | hypothetical protein |
| 0.0 | 0.0 | 0.0 | 1863 | 76 | hypothetical protein |
| 0.0 | 0.0 | 0.0 | 1864 | 47 | hypothetical protein |
| 0.0 | 0.0 | 0.0 | 1865 | 65 | hypothetical protein |
| 0.0 | 0.0 | 0.0 | 1866 | 56 | hypothetical protein |
| 0.0 | 0.0 | 0.0 | 1867 | 88 | hypothetical protein |
| 0.0 | 0.0 | 0.0 | 1868 | 77 | hypothetical protein |
| 0.0 | 0.0 | 0.0 | 1869 | 157 | hypothetical protein |
| 0.0 | 0.0 | 0.0 | 1870 | 99 | hypothetical protein |
| 0.0 | 0.0 | 0.0 | 1871 | 186 | hypothetical protein |
| 0.0 | 0.0 | 0.0 | 1872 | 54 | hypothetical protein |
| 0.0 | 0.0 | 0.0 | 1916 | 52 | Adhesin of unknown specificity SdrC |

| % Identity | | | 1519 | | |
|---|---|---|---|---|---|
| ED98 | Ch21 | B4-59C | PEG | Length | function |
| 0.0 | 0.0 | 0.0 | 1917 | 205 | Type I restriction-modification system, DNA-methyltransferase subunit M / Type I restriction-modification system, specificity subunit S; or LPXT- cell wall anchor domain, and/or fibrinogen-binding protein |
| 24.3 | 24.3 | 24.3 | 1919 | 178 | Aminoglycoside N6'-acetyltransferase |
| 0.0 | 0.0 | 0.0 | 1921 | 144 | hypothetical protein |
| 31.8 | 31.8 | 31.8 | 1922 | 110 | Transcriptional regulator, ArsR family |
| 45.0 | 0.0 | 0.0 | 1923 | 173 | Plasmid recombination, MobE mobilization protein |
| 0.0 | 0.0 | 100.0 | 2179 | 95 | Hypothetical protein in SaPI |
| 30.1 | 30.1 | 100.0 | 2180 | 121 | Hypothetical protein in SaPI |
| 0.0 | 0.0 | 100.0 | 2181 | 486 | Virulence-associated protein E in SaPI |
| 0.0 | 0.0 | 100.0 | 2189 | 45 | hypothetical protein |
| 29.4 | 29.4 | 100.0 | 2191 | 193 | hypothetical protein |
| 0.0 | 0.0 | 100.0 | 2192 | 59 | hypothetical protein |
| 0.0 | 0.0 | 0.0 | 2358 | 280 | hypothetical protein |
| 0.0 | 0.0 | 0.0 | 2369 | 205 | hypothetical protein |
| 0.0 | 0.0 | 0.0 | 2384 | 146 | Phage holin |
| 100.0 | 100.0 | 0.0 | 2688 | 39 | hypothetical protein |
